# *Nr5a2* is essential for morula development

**DOI:** 10.1101/2023.01.16.524255

**Authors:** Nicola Festuccia, Sandrine Vandormael-Pournin, Almira Chervova, Anna Geiselman, Francina Langa-Vives, Rémi-Xavier Coux, Inma Gonzalez, Michel Cohen-Tannoudji, Pablo Navarro

## Abstract

Early embryogenesis is driven by transcription factors (TFs) that first activate the zygotic genome and then specify the lineages constituting the blastocyst. While the TFs specifying the blastocyst’s lineages are well characterised, those playing earlier roles are ill-defined. Using mouse models of the TF *Nr5a2*, we show that *Nr5a2*^*-/-*^ embryos arrest at the early morula stage and exhibit overt phenotypical problems such as altered lineage specification, frequent mitotic failure and substantial chromosome segregation defects. Transcriptomic profiling shows that NR5A2 is a master regulator required for appropriate expression of thousands of genes at the 8-cells stage, including lineage-specifying TFs and genes involved in mitosis, telomere maintenance and DNA repair. We conclude that NR5A2 coordinates proliferation, genome stability and lineage specification to ensure proper morula development.

## Introduction

In mammals, preimplantation development corresponds to the transition from a single cell, the zygote, to a complex structure, the blastocyst, constituted of three lineages that will subsequently be the founders of all extra-embryonic and embryonic tissues. The specification of the lineages of the blastocyst starts in the morula, the ensemble of 16 to 32 cells resulting from the successive cleavages of the zygote and compaction, and is mediated by the hierarchical action of several transcription factors (TFs): TEAD4, GATA3 and CDX2 or SOX2 and OCT4, among others, will first specify the extra-embryonic trophectoderm (TE) and the inner cell mass (ICM), forming early blastocysts; NANOG and GATA6 will subsequently segregate the embryonic pluripotent epiblast (EPI) or the extra-embryonic primitive endoderm (PrE), forming late blastocysts competent for implantation and further development **(1)**. While these and other TFs involved in the formation of the blastocyst have been extensively studied, how the zygote progresses to the morula stage remains unclear.

TFs have also been associated with the first regulatory step starting mouse development, the zygotic genome activation (ZGA), which jumpstarts transcription at the 2-cells (2C) stage **(2)**. In contrast to lineage determinants, many of which perform essential roles in the formation of the blastocyst, the invalidation of TFs involved in ZGA, including DUX, RARG, DPPA2/4, NFYA and YAP1, typically does not compromise ZGA or progression beyond the 2C stage **(3-10)**. Recently, however, the TF NR5A2 has been suggested to play a preponderant role during ZGA: upon chemical inhibition of NR5A2, developmental arrest was observed from the 2C stage **(11)**. Even though *Nr5a2* knock-out models need to be developed to formally establish its essential role during ZGA, specific TFs have been identified as regulators of the early (ZGA) and late events (lineage specification) taking place during preimplantation development. Whether any TF is required to enable progression from the 2C stage to a functional morula from where early lineages will emerge remains unknown.

The potential involvement of NR5A2, in combination with ESRRB (two related nuclear receptors), in preimplantation development is of interest given the established role of these two TFs in the control of pluripotency in mouse Embryonic Stem (ES) cells **(12)**. Indeed, they conjunctly control thousands of regulatory elements, in interphase but also during mitosis **(13)**, to confer robustness to the expression and activity of major regulators of pluripotency such as OCT4, SOX2 and NANOG. Hence, NR5A2 and ESRRB could represent global regulators of early mouse embryogenesis, controlling events linking ZGA to the formation of the blastocyst. In this study, we aimed at addressing this hypothesis by using gene knock-outs of *Nr5a2* and *Esrrb*. We found that neither NR5A2 nor ESRRB are required, alone or in combination, to execute ZGA and progress beyond the 2C embryo. Instead, we found that NR5A2 alone is absolutely essential for the development of a mature morula capable of initiating lineage specification and developing into a blastocyst.

## Results

### Maternally expressed NR5A2 and ESRRB are dispensable for embryogenesis

To study the roles of NR5A2 and ESRRB during preimplantation development we generated female mice carrying conditional knock-out alleles for *Esrrb* **(14)** and/or *Nr5a2* **(15)**, in addition to *Zp3:Cre* **(16)**. Allele recombination during oocyte growth allowed testing whether development proceeds in the complete absence of maternal (mKO), zygotic (zKO) or maternal and zygotic (mzKO) expression of either or both genes (see methods, and **Table S1**). NR5A2, with support from ESRRB, has been recently reported to function as a major regulator of ZGA **(11)**, implying that ablating the maternal contribution of these genes should trigger defects of functional consequence. We thus first sought to examine the effects of inducing the loss of maternal *Nr5a2/Esrrb*. Contrary to expectations, females generating oocytes devoid of NR5A2, ESRRB, or both TFs, gave birth to litters of equal size to matched controls, efficiently transmitting the deleted alleles **(Fig 1A, Table S1)**. This rules out the possibility that the maternal inheritance of NR5A2 or ESRRB performs a determinant function. In accordance, low NR5A2 expression was observed before or around the time of ZGA **(Fig 1B, S1A)** compared to later stages, as controlled by knock-out embryos and confirmed by Ribo-seq **(Fig S1B) (17)**, and RNA-seq **(Fig S1C) (18)**. Similarly, Esrrb-TdTomato reporter mice revealed that ESRRB is not maternally inherited at significant levels **(Fig 1C, S1D)**. Taken together, these results suggest that *Nr5a2* and *Esrrb* are unlikely to represent maternal regulators of ZGA; they may however exert a developmental function after ZGA allows accumulation of their mRNAs and proteins.

**Fig. 1.**
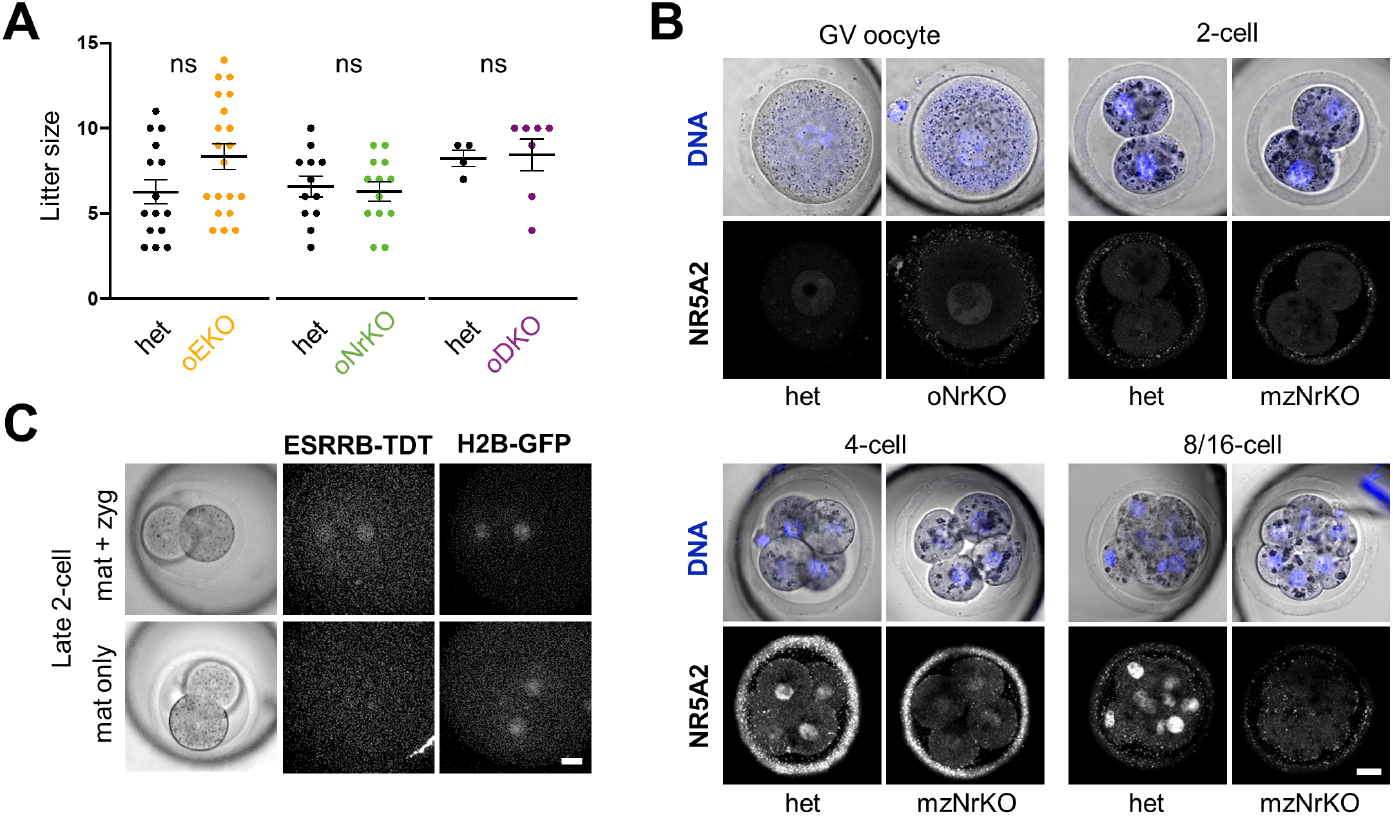
Preimplantation development does not require maternal NR5A2 and ESRRB. **(A)** Size of litters from crosses between females generating oocytes devoid of NR5A2 and/or ESRRB (oEKO: *Zp3Cre*^*tg/0*^; *Esrrb*^*flox/del*^, oNrKO: *Zp3Cre*^*tg/0*^; *Nr5a2*^*flox/del*^, oDKO: *Zp3Cre*^*tg/0*^; *Esrrb*^*flox/del*^; *Nr5a2*^*flox/del*^) and wild-type males. Littermate females (*Zp3Cre*^*tg/0*^; *Esrrb*^*flox/+*^ and/or *Nr5a2*^*flox/+*^) were used as controls. **(B)** Immunofluorescence of NR5A2 in germinal vesicle (GV) and early embryos. Stainings of oNrKO GV and mzNrKO embryos show a low nuclear immunostaining background mainly in GV oocytes. Scale bars equal 20 um in all panels. **(C)** Expression of ESRRB-TdT fusion protein in late 2C embryos obtained from a cross between an *EsrrbTdT* knock-in female with a *H2b-Egfp* transgenic male. Upper panels show an embryo having inherited both the *EsrrbTdT* knock-in allele and any eventual maternal mRNAs and protein store (mat +zyg), while the embryo shown in lower panels did not inherit the knock-in allele (mat only).

### Zygotically expressed NR5A2 is essential for the development of a viable morula

We next sought to assess the effects of ablating maternal and zygotic expression of the two TFs (mzKO). The individual genetic disruption of *Esrrb* and *Nr5a2* is known to trigger developmental defects only around/after implantation, in extra-embryonic and embryonic lineages respectively **(19-22)**. We thus anticipated any eventual earlier phenotype to manifest exclusively in double knock-out (DKO) embryos. In agreement, *Esrrb* mzKO was inconsequential at the stages analysed, and morphologically normal mutant blastocysts having completed lineage segregation could be recovered at E4.25 **(Fig 2A, Table S1)**. In contrast, and unexpectedly, all E3.5-E4.25 *Nr5a2* mzKO embryos showed profound morphological abnormalities and a marked developmental delay **(Fig 2B,C, Table S1)**, accompanied by reduced cell numbers, degenerated nuclear morphology, and evident signs of disordered progression through cell division **(Fig 2D)**. To exclude potential biases due to the genetic background (prevalently C57BL/6) and assess the contribution of zygotic NR5A2 expression, we analysed *Nr5a2* zKO generated on a mixed C57BL/6;CD1 background, which confirmed the essential role of *Nr5a2* **(Fig S2A,B)**. Ex-vivo culture of 4C embryos revealed that developmental delay/defects exacerbate progressively after the 8C stage, and by the time control embryos have formed blastocysts, mutant morulae degenerate **(Fig 2E, S2D)**. At E2.75 (8-16C), expression of GATA3 and NANOG but not of GATA6, key TFs of TE, EPI and PrE respectively, was radically affected in both *Nr5a2* mzKO and mzDKO embryos **(Fig 2F, S2A)**. At E2.5, mutants already showed a clear developmental delay **(Fig 2C)** and incipient NANOG expression was compromised **(Fig 2G)**. Other pluripotency TFs seemed either less (SOX2) or hardly (OCT4) affected **(Fig S2C)**. Therefore, NR5A2 is essential for the formation of a viable morula.

**Fig. 2.**
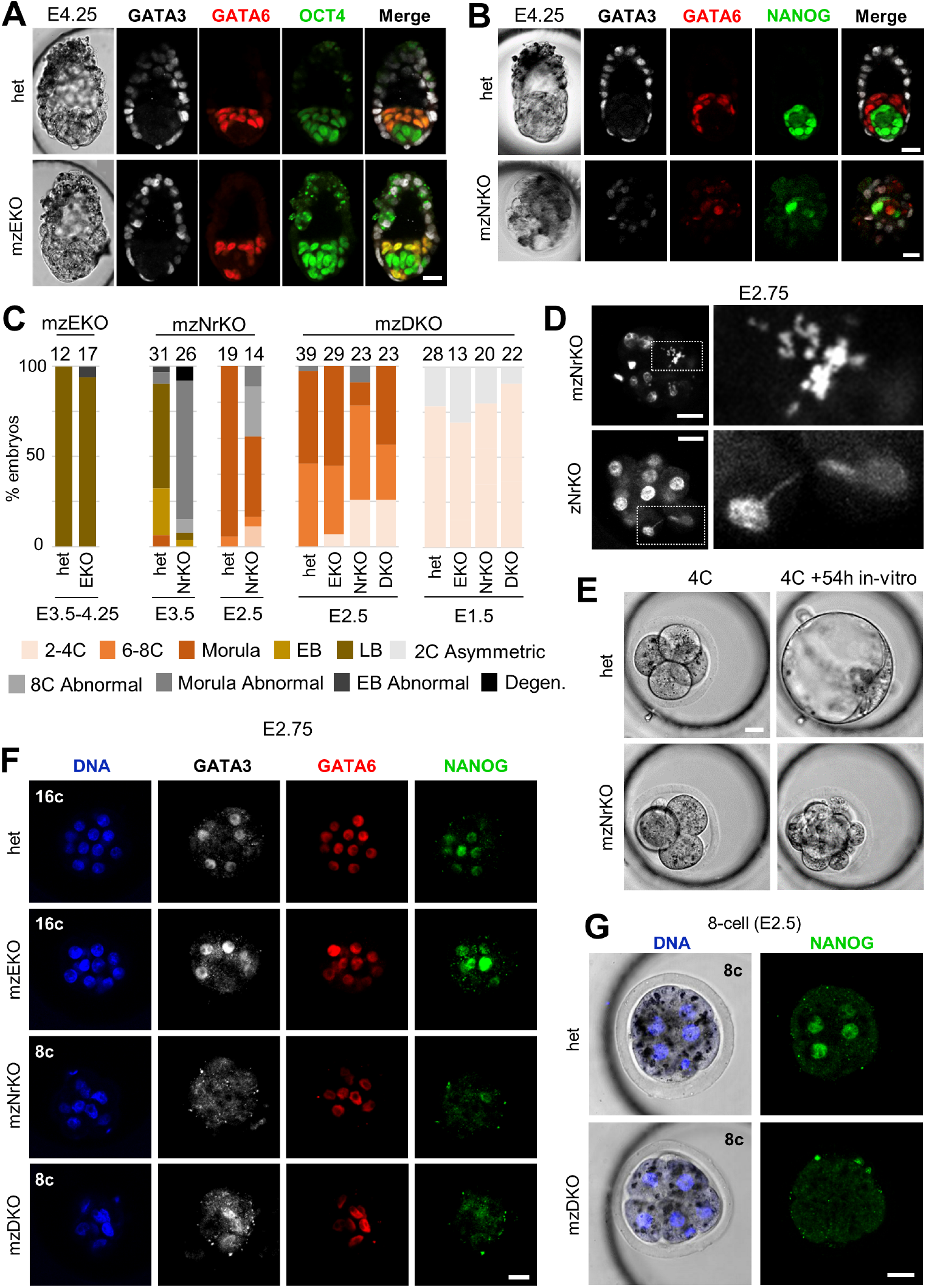
*Nr5a2* is required for the formation of a viable morula. **(A)** GATA3, GATA6 and OCT4 immunostaining of maternal and zygotic *Esrrb* heterozygous and KO embryos at E4.25. Scale bars equal 20 um in all panels. **(B)** GATA3, GATA6 and NANOG immunostaining of maternal and zygotic *Nr5a2* heterozygous and KO embryos at E4.25. **(C)** Rate of development stages observed for each genotype in E1.5 to E4.25 litters from crosses generating mzEKO, mzNrKO and mzDKO embryos. Degen: Degenerated. **(D)** Hoechst DNA staining highlights mitotic defects (misaligned chromosomes at metaphase (up) and chromatin bridges (down)) in zygotic or maternal and zygotic *Nr5a2* KO embryos collected at E2.75. **(E)** Maternal and zygotic *Nr5a2* heterozygous and KO embryos dissected at the 4C stage (52h post-hCG) and cultured for 54 hours in-vitro. **(F)** GATA3, GATA6 and NANOG immunostaining of mzDHet, mzEKO, mzNrKO and mzDKO littermate embryos at E2.75. **(G)** NANOG immunostaining of mzDHet and mzDKO littermate 8-cell (E2.5) embryos.

### *Nr5a2* KO results in global transcriptional deregulation only at the 8C-stage

Interfering with NR5A2 activity in cultured zygotes has profound consequences on ZGA and results in developmental blockade or delay starting from the 2C stage **(11)**. However, the complete genetic loss of *Nr5a2* triggers major defects only after the 8C stage. Therefore, we charted the transcriptional response to the mzKO of *Nr5a2, Esrrb* or both TFs in-utero, immediately after ZGA in late 2C embryos, as well as in 4C and 8C embryos. Flash-seq based **(23)** analysis on 122 embryos distributed across 4 genotypes **(Table S2)** determined that in the absence of the two TFs (mzDKO) only 407 genes are affected immediately after ZGA (152 up, 255 down), a number that modestly increases at the 4C stage (358 up, 428 down). In contrast, at the 8C stage, concomitantly with the onset of cellular and morphological defects, nearly 3000 genes were found deregulated (1713 up, 1253 down; **Fig 3A-C**). The observed changes were validated by absolute transcript counts **(Fig S3A)**. Gene expression trends at the 2 and 4C stage were found largely correlated **(Fig 3A)**, with a prominent overlap between differentially expressed genes (DEGs) that fail to be expressed in mutant embryos (DOWN, **Fig 3B**). Therefore, while NR5A2 and ESRRB are not strictly required to execute ZGA, this analysis confirms that they display an early role in gene regulation. Concordant regulation is also observed at the 8C stage, but diluted by the high number of genes that start being affected only at this time. In line with single KO phenotypes, *Esrrb* KO was found quasi-inconsequential at the 8C stage whereas *Nr5a2* KO recapitulated the effect of the DKO **(Fig 3C, S3B)**. Nevertheless, a modest but clear effect of ESRRB was observed at the 2C stage, where *Esrrb* KO and DKO DEGs appeared concordantly regulated. Moreover, matching the delay observed in mutant embryos, Principal Component Analysis of single embryos delineated a developmental trajectory on which *Nr5a2* KO and DKO embryos start diverging from wild-type and *Esrrb* KO embryos only at the 4C stage and clearly segregate from controls at the 8C stage **(Fig 3D)**. This suggests that when incipient expression of both TFs is low in late 2C embryos, ESRRB and NR5A2 might cooperatively control a limited set of genes. Later, at the 8C stage, when NR5A2 expression peaks **(Fig 1B, S1A)**, this TF acquires a dominant and broader role. Finally, to further establish NR5A2 as a major direct regulator of the 8C transcriptome we performed de-novo motif discovery over accessible regions surrounding NR5A2-responsive genes in 2, 4 and 8C embryos **(Fig 3E, Table S3)**. This analysis revealed the enrichment of a variant of the sequence bound by orphan nuclear receptors (TCAAGG(T/C)CA), that we previously identified as functionally determinant in ES cells **(12)**, and which is nearly identical to the known NR5A2 binding consensus. Altogether, our results indicate that NR5A2 is a major direct regulator of hundreds of genes at the 8C stage, a time at which the phenotypical consequences of its genetic invalidation start manifesting.

**Fig. 3.**
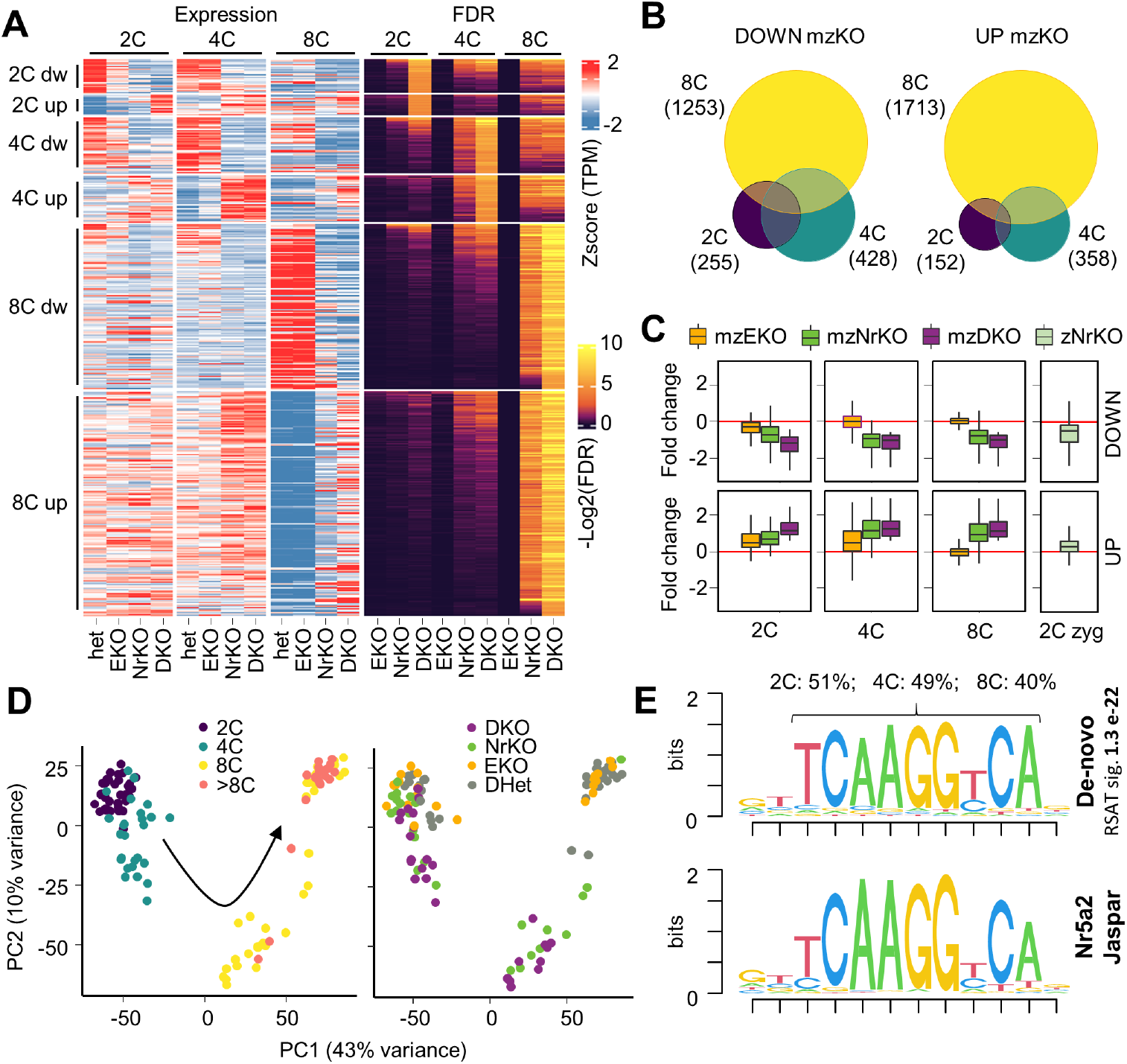
Transcription is progressively deregulated in *Nr5a2* KO embryos, from the 2C to the 8C stage. **(A)** Heatmap showing relative expression (Z-scores) of genes significantly affected - absolute Fold change >1.5; False discovery rate (FDR) < 0.1; cutoff 10 transcripts per million (TPM) - after the loss of maternal and zygotic expression of *Esrrb, Nr5a2* or both genes at the 2, 4 or 8-cell stage. **(B)** Number and overlap between the genes differentially expressed (as in A) in response to the loss of *Esrrb, Nr5a2* or both genes at different stages of development. **(C)** Fold change in expression of genes significantly affected (as in A) in response to the maternal and zygotic KO of *Nr5a2* and/or *Esrrb*. The fold change of the same set of genes in response to the zygotic knockout of *Nr5a2* at the 2C-stage is shown alongside (2C zyg). **(D)** Principal component analysis of the transcriptome from individual embryos color coded to highlight differences in genotypes (right) or developmental stage (left). The black arrow intends to highlight an ideal progression along a pseudo-time of development. **(E)** ESRRB/NR5A2 binding sequence identified by de novo TF binding motif discovery over accessible regions surrounding the NR5A2-responsive genes in 2, 4 and 8C embryos, and NR5A2 binding consensus retrieved from the Jaspar database. The percentage of responsive genes (as in A) associated with an accessible region containing the NR5A2/ESRRB motif is reported at the top.

### ZGA proceeds in the complete absence of NR5A2

In order to refine our understanding of the role of NR5A2 and ESRRB in ZGA, we analysed the effect of the complete absence of these TFs (mzDKO) on a set of 2000 genes previously shown to be robustly induced during ZGA **(Fig S4A) (11)**. We observed a limited effect of NR5A2 and ESRRB on these genes at both late 2C (23 up, 74 down) and 4C (68 up, 117 down) stages **(Fig 4A left)**. These results indicate that gene activation is largely normal during ZGA in the absence of NR5A2. Moreover, the small effects observed could be vastly attributed to the function of zygotically expressed NR5A2, as equivalent changes were observed after its exclusively zygotic ablation **(Fig 4A right)**. Next, we asked whether NR5A2 might instead have an effect on the expression of genes that, as many known regulators of ZGA, peak at the 2C stage (early/mid/late 2C; 226 genes, **Fig S4A**), including *Zscan4b-f, Duxf3*, and *Tcstv1/3*. Expression of these 2C-specific transcripts was not affected in mutants at the 2C stage. Instead, these genes maintained heterogeneously increased expression levels until the 8C stage **(Fig 4B)**. This may be a consequence of the developmental delay observed in mutant embryos **(Fig 2C,E and 3D)** or may hint at a specific role of NR5A2 in facilitating the extinction of the 2C state. We then turned our attention to repetitive elements, since specific groups, Mouse Endogenous Retrovirus type-L (MERVLs)/non-autonomous Mammalian apparent LTR Retrotransposons (MaLRs), promote expression of proximal genes during ZGA **(24**,**25)**. Moreover, NR5A2 binds SINE elements, in particular B1/Alu types, with similar functions during ZGA **(11**,**26**,**27)**. However, no element belonging to either class showed significant changes in mutant embryos at the 2 or 4C stage **(Fig S4B)**. Similarly to 2C-specific single-copy genes, improperly elevated levels of specific MERVL/MaLR transcripts were detected in 8C embryos **(Fig 4B)**, as also observed in ES cells **(Fig S4C) (28)**, supporting the possibility that NR5A2 prevents re-expression of 2C-specific transcripts as development proceeds. SINE elements, which remain expressed at 8C, were instead down-regulated **(Fig 4C)**, confirming the positive regulation of this class of repeats by NR5A2 **(11**,**26)**. We note that, although not passing significance filters, a global reduction in expression was noticeable for SINE and MERVL/MaLR elements in 2C mutant embryos **(Fig S4B)**. It is thus likely that zygotically expressed NR5A2 contributes to the regulation of gene expression immediately after the beginning of ZGA and afterwards, by binding to or in proximity of these types of repeats. In agreement with this view, and as reported **(11)**, motif analyses at putative regulatory regions **(24)** of NR5A2-responsive genes at 2C stage revealed enrichment for a slightly degenerated and longer version of the NR5A2 motif, possibly composed of a direct repeat of two nuclear receptor binding elements, matching a fragment of the consensus sequence of the YR5 CA variant of SINE elements **(11) (Fig 4D, Table S3)**. Our results indicate that NR5A2 is not an essential regulator of ZGA even though it regulates a small subset of genes at the 2C stage, possibly by binding SINEs.

**Fig. 4.**
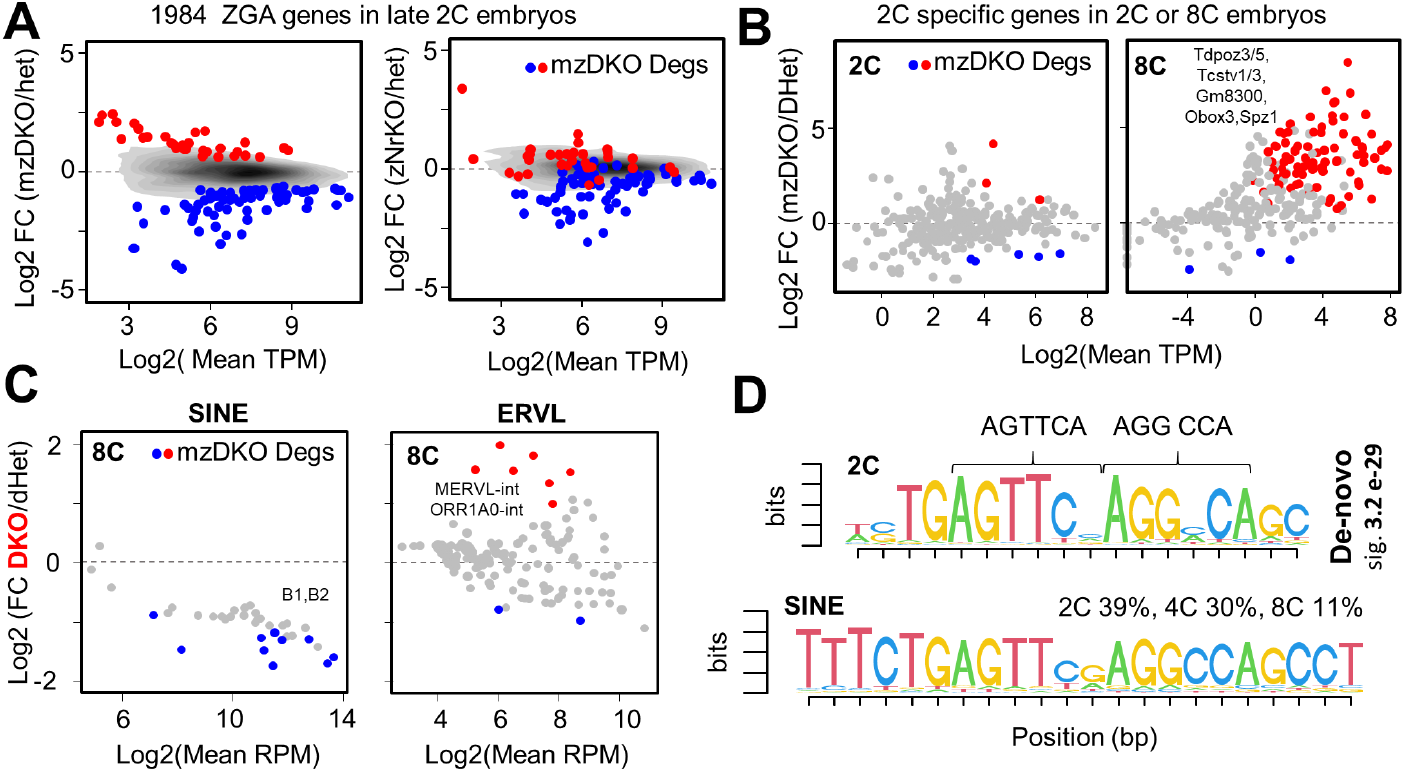
Zygotic genome activation proceeds with minor defects in the complete absence of *Nr5a2/Esrrb*. **(A)** Fold change in expression of 1984 genes activated during ZGA (as defined previously **(11)**, and expressed in our datasets) in response to the maternal and zygotic KO of *Nr5a2* and *Esrrb*, or the zygotic KO of *Nr5a2* alone at the late 2C stage. Red and blue dots identify significantly upregulated or downregulated genes - absolute Fold change > 1.5; False discovery rate (FDR) < 0.1; cutoff 10 transcripts per million (TPM) - in mzDKO embryos. The gray shades represent the density distribution of all genes, divided in 10 bins. **(B)** Fold change in expression of 290 genes peaking at the 2C stage (see methods) in response to the maternal and zygotic KO of *Nr5a2* and *Esrrb*, at the late 2C or 8C stage. Red and blue identify significantly upregulated or downregulated genes in mzDKO embryos at the respective stage (as in A), gray all other expressed genes. **(C)** Fold change in expression of SINE or ERVL repeats in response to the maternal and zygotic KO of *Nr5a2* and *Esrrb*, at the 8C stage. Red and blue identify significantly upregulated or downregulated repeats in mzDKO embryos (absolute Fold change > 1.5; False discovery rate (FDR) < 0.1). **(D)** ESRRB/NR5A2 binding sequence identified by de novo TF binding motif discovery over accessible regions surrounding the NR5A2 responsive genes in 2C embryos only, and a fragment of the SINE Y5 variant consensus sequence.

### *Nr5a2* controls genes involved in lineage specification, cell division and genomic stability

To explore the molecular causes underlying the defects observed in the absence of *Nr5a2*, we performed gene ontology analyses after combining all differentially expressed genes throughout experimental conditions **(Table S4)**. Clustering the first 150 most enriched terms revealed several groups of related ontologies that directly echoed the observed phenotypes **(Fig 5A)**. A prominent association with developmental terms was evident, such as blastocyst and placenta development (Groups 2 and 6), characterized by major key effectors of lineage specification into either the EPI (*Nanog, Klf4*, and *Sox15*) or the TE (*Tead4, Tfap2c, Gata3, Klf5* and *Cdx2*) that are down-regulated in *Nr5a2* KO and dKO embryos mostly at the 8C stage **(Fig 5B)**. Some exceptions exist, such as genes with an early developmental function (*Sox15, Tead4*) **(29**,**30)** that are downregulated from the 2C stage, and PrE genes that are upregulated (*Gata4, Sox17, Pdgfra*; **Fig 5B**). Thus, NR5A2 appears as an upstream regulator of many of the most relevant TFs driving the formation of the blastocyst. Another cluster of gene ontology terms (Group 1) was directly related to the mitotic and cell division defects observed in *Nr5a2* KO and dKO embryos **(Fig 5A)**. This cluster included genes important for the mitotic regulation of the centromeres (*Cenpa, Haspin, Borealin* and *Shugosin*) or more general regulators of the cell cycle (members of the APC/C complex – *Cul3* and *Ube2c* –, the mitotic *Cdk1* kinase and its regulators *Cdc25* and *Mcph1*, or the G1/S regulator *Cyclin E*). Many of these genes were downregulated from the 2C or 4C stage **(Fig 5B)**, likely underlying the impaired ability of mutant blastomeres to proliferate and properly partition the genetic material during mitosis. Finally, a last cluster of genes associated with DNA repair, homologous recombination and telomere maintenance (Group 7) might also be related to the low cell numbers observed in *Nr5a2* KO and dKO embryos **(Fig 5A)**. These include telomeric proteins and genes involved in DNA replication and repair that have a central function in telomere maintenance, preferentially by the ALT pathway **(31)**, which contributes to telomere lengthening in early embryos **(32)**. Notably, nuclear receptors play an active role in this pathway by binding to variant telomeric repeats in cancer cells **(33)**. Hence the deregulation of this set of genes **(Fig 5B)** might be part of a broader response to telomere damage in the absence of NR5A2, which directly binds to subtelomeric regions in ES cells **(Fig 5C**,**D)**. In addition to its role as an upstream regulator of lineage drivers and cell cycle genes, NR5A2 may thus play a direct role in telomere homeostasis during preimplantation development.

**Fig. 5.**
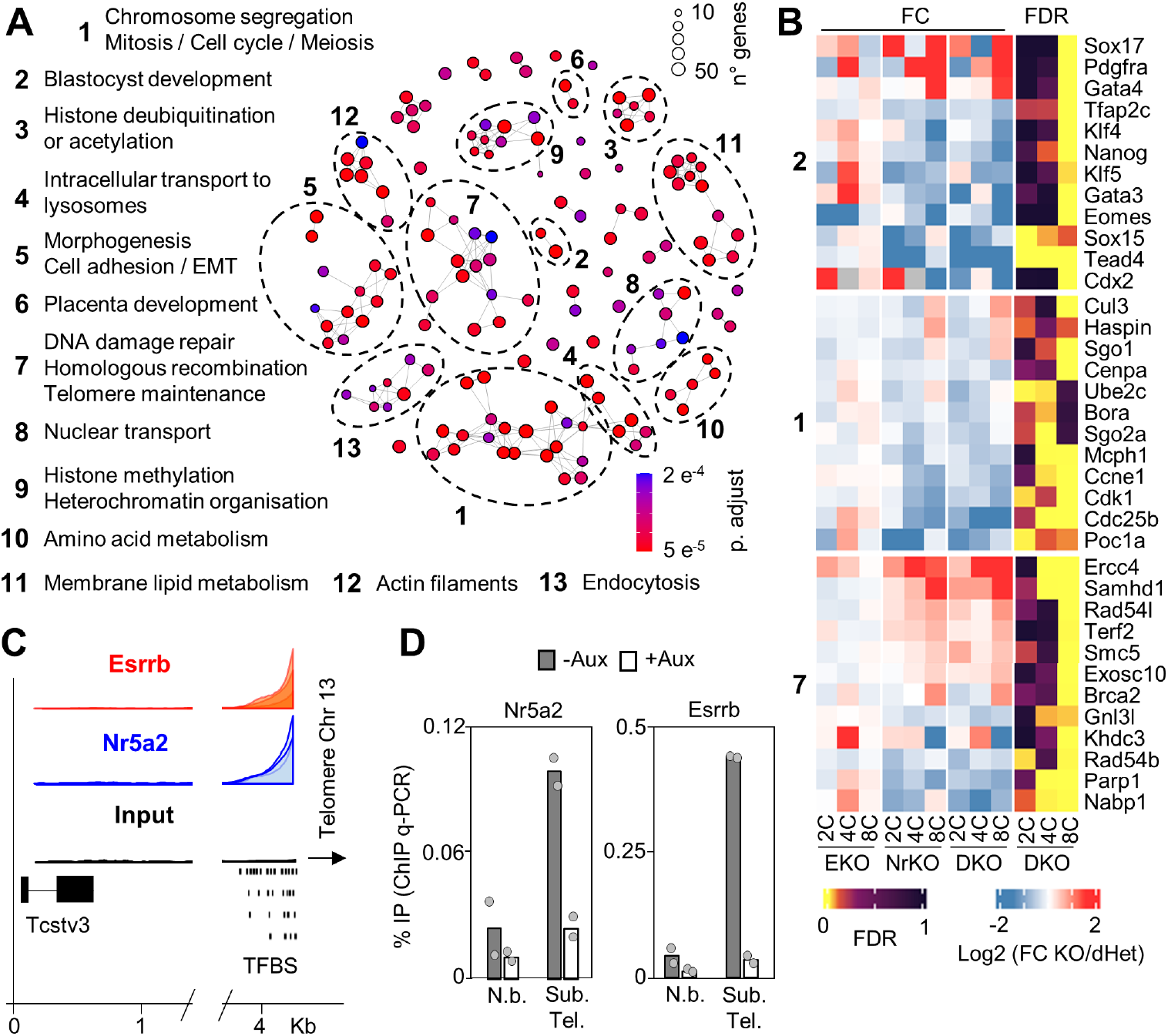
NR5A2 controls master lineage TFs and genes involved in DNA repair, telomere maintenance and cell division. **(A)** Map showing the relation between gene ontology terms enriched among the genes that respond to the loss of NR5A2 and/or ESRRB in 2, 4 and 8C embryos. Terms sharing a significant number of associated genes are linked by gray lines and clustered together. The number of responsive genes associated to each term is indicated by the circle size, the adjusted p-value for the enrichment of each term by the colorscale. Numbering is based on the p-value of the most enriched term in each cluster. **(B)** Heatmap showing the fold change in expression of genes significantly affected - absolute Fold change > 1.5; False discovery rate (FDR) < 0.1; cutoff 5 transcripts per million (TPM) - after the loss of maternal and zygotic expression of ERRB, NR5A2 or both genes at the 2, 4 or 8-cell stage. The FDR for the differential expression of each gene is plotted alongside. **(C)** NR5A2 and ESRRB binding in ES cells **(12**,**13)** in proximity of the subtelomeric region near *Tcstv3* on Chr13. Note the presence of clusters of degenerate NR5A2/ESRRB binding sites. Tracks from the input are shown in support of the specificity of the signal observed. **(D)** Nr5a2 and Esrrb binding at the same sub-telomeric regions as in C, assessed by ChIP q-PCR in ES cells in which NR5A2 and ESRRB can be depleted by addition of Auxin **(13)**. N.b.: Non binding

## Discussion

In this study, we document the first example of a developmental TF that becomes active soon after the beginning of ZGA and shows a progressively increasing importance in coordinating basic functions such as proliferation, the maintenance of genomic stability and the control of the transcriptional events that set the ground for the onset of lineage specification. Our study therefore fills a hereto missing link, identifying a TF, *Nr5a2*, that is able to shepherd development of the embryo from the period of ZGA to the moment in which the first cell fate specification events occur in the morula.

We show that *Nr5a2*^*-/-*^ embryos arrest at the early morula stage; however, previous studies reported a later phenotype **(19**,**20**,**22)**. While the basis of this discrepancy is unclear but might be linked to different genetic backgrounds (129Sv-MF1 or not reported in previous studies and C57BL/6 or C57BL/6;CD1 here), independent work has also shown profound detrimental effects on preimplantation development upon *Nr5a2* knock-down, inhibition with chemical inhibitors **(11)** and gene editing in zygotes **(26)**. Together with our work, these studies establish NR5A2 as a key TF for early mouse embryogenesis. Recently, it has been suggested that inhibiting NR5A2 activity affects expression of a vast number of genes during ZGA, leading to an extremely early embryonic arrest **(11)**. Our analyses based on the genetic inactivation of maternal and/or zygotic expression of *Nr5a2* excludes such an essential role. However, we show that the zygotic expression of NR5A2 does indeed contribute to ZGA, as confirmed by *Nr5a2* editing in zygotes **(26)**. In light of these observations, the results obtained by the use of chemical inhibitors are particularly interesting. While directed against NR5A2, SR1848 strongly inhibits NR2C2 **(11)** and possibly more nuclear receptors expressed at the 2C stage **(26)**. Given that these TFs often share the same DNA binding motif, and that we have shown a high redundancy in their chromatin association **(12**,**13)**, it is possible that this class of TFs represents, as a whole, the first example of a so far elusive sequence specific activity essential to jumpstart zygotic transcription in mammals. In this regard, it is noteworthy that the motif identified at genes responding to NR5A2 in 2C stage embryos, which matches SINE B1/Alu repeats, is constituted of a direct tandem of two basic nuclear receptor response elements. This setup promotes heterodimerization between nuclear receptors **(34)**, as shown for RXR-RAR dimers **(35)**, hinting at an interplay between NR5A2 and these or other nuclear receptors in the recognition of SINE elements and the promotion of ZGA. More generally, and as illustrated by *Dppa2* **(9)**, *Klf5* **(36)** or *Dux* **(37)**, our results on *Nr5a2* highlight that many sequence specific TFs, while not maternally inherited and themselves upregulated during ZGA, engage in a reinforcing feedback loop that boosts gene activation. Nonetheless, none of these factor appears individually essential.

Besides the involvement in ZGA, our study identifies a distinct and important role for *Nr5a2* as the first example of a single TF essential for the formation of a viable morula. This is of crucial importance: while the players controlling lineage determination have been extensively characterised, which factor prepare the embryo for the beginning of these cell fate choices remains poorly understood. Unsurprisingly, inactivation of TFs controlling TE fate, such as *Tead4* **(30**,**38)** and *Cdx2* **(39**,**40)**, leads to the closest phenotypes to that observed in *Nr5a2* mutants. Given the fact that expression of *Tead4* and *Cdx2* is severely altered in *Nr5a2* KO embryos, it is possible that these genes might be important mediators of the lineage specification phenotype of *Nr5a2*^*-/-*^ embryos. However, NR5A2 displays a dual activation function of both TE and EPI drivers and should therefore be seen as a TF promoting developmental bi-potential, as shown for *Klf5* **(36)**, a prime NR5A2 target whose inactivation leads to pre-implantation arrest at later stages **(36**,**41)**. Finally, NR5A2 appears to be unique in its extraordinarily early requirement for proliferation and genomic stability during cleavage stages. This activity might be exerted either indirectly, by controlling genes important to assemble the mitotic machinery or repair DNA, or directly, by targeting degenerate subtelomeric repeats and controlling the ALT pathway. Strikingly, such telomeric association was originally shown for NR2C2 **(33)**, which is also expressed soon after ZGA and can be inhibited with SR1848, like NR5A2 **(11**,**26)**. The role of orphan receptors in telomere maintenance and their involvement in the ALT pathway may therefore go beyond the context of cancer and be of primarily importance during preimplantation development.

Overall, nuclear receptors might be the class of TF showing the earliest essential role during development, starting at the 8C stage. This is precisely the period when a second major wave of transcription, named mid-preimplantation gene activation (MGA), has been proposed to take place **(42)**. Indeed, gene expression steadily increases after ZGA, with expression of many genes peaking only at 4-8C, when conventional features of the transcriptional cycle are also gradually established **(43)**. It will therefore be important to test whether NR5A2 and other nuclear receptors participate in establishing these mature traits of gene regulation, essential to effectively direct the first cell fate specification events.

## Acknowledgements

We acknowledge the Mouse Genetics Engineering Center, Biomics, Photonic BioImaging and Image Analysis Hub platforms of Institut Pasteur. We also acknowledge Diana Voicu and Michael Imbeault for critical reading of the manuscript and for discussions on repetitive elements and their analyses. This study was supported by recurrent funding from Institut Pasteur (PN), CNRS (NF, MCT, PN), Revive Investissement d’Avenir ANR-10-LABX-73 (MCT and PN) and by European Research Council grant ERC-CoG-2017 BIND (PN).

## Author contributions

NF performed gene expression, ChIP, and most computational analyses. Embryo collection, culture, and processing was carried out by SVP. AC analysed FLASH-seq data. AG contributed preparatory embryo immunostainings. IG, RXC and FLV derived ESRRB-TDT reporter mice. DV and ME assisted with analyses of DNA repeats. NF conceived the project, with the contribution of MCT. NF, MCT and PN supervised the work, analysed the data and wrote the manuscript.

## Supplementary information

Five supplementary figures can be found at the end of this document. Four Supplementary Tables are available online. Supplementary Methods can be found online.

---

**Supplementary Information, Fig.S 1.**
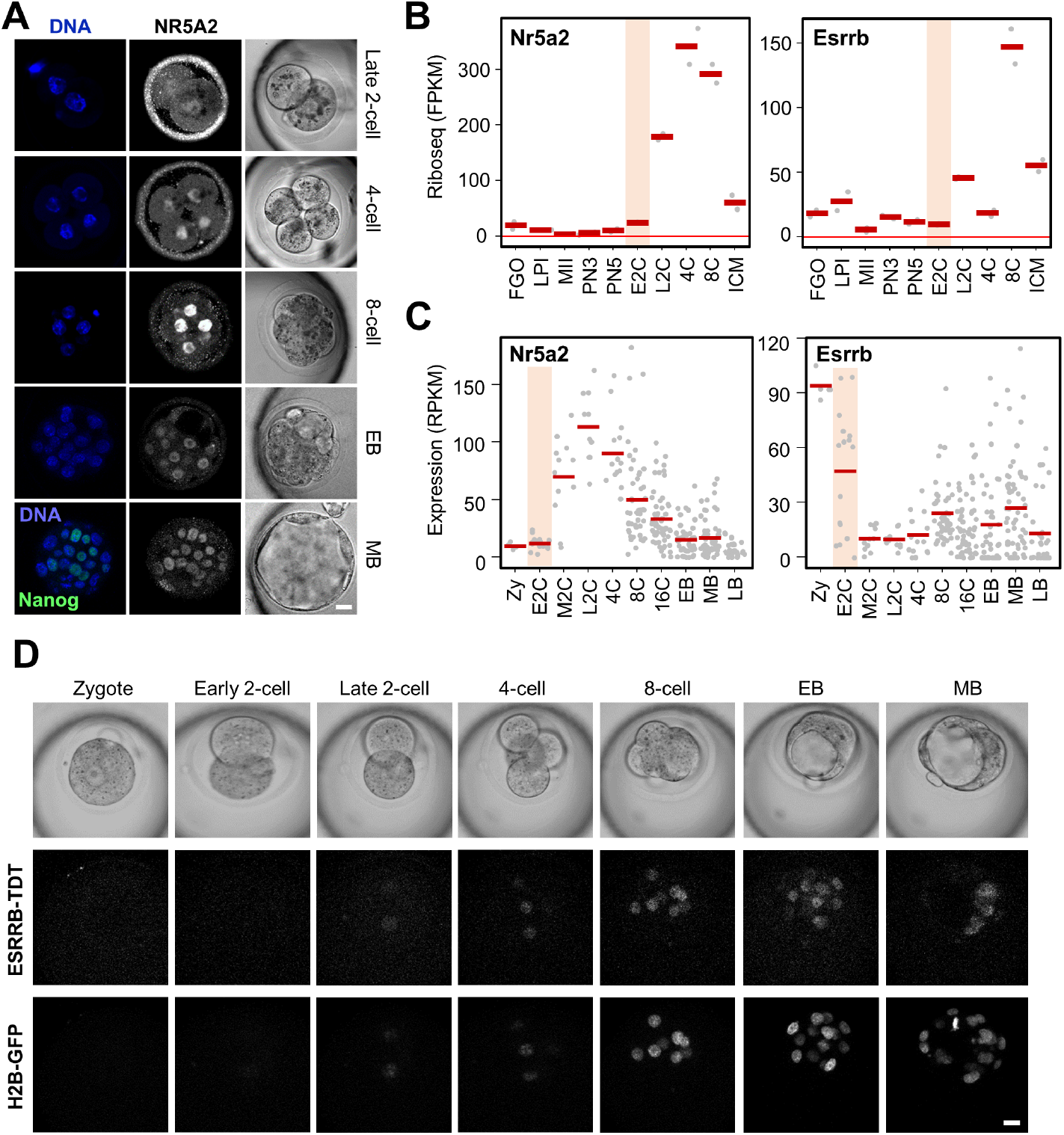
No or marginal levels of NR5A2 and ESRRB maternal stores. (A) NR5A2 immunofluorescent staining in wild-type embryos at different stages of development. (B,C) Levels of mRNA undergoing translation **(17)**, or total mRNA **(18)**, in oocytes or embryos/single blastomeres at different developmental stages. Zy: Zygote; FGO: Fully grown oocyte; LPI: late prometaphase I oocytes; MII: MII oocytes; PN3/5: Zygote at pronuclear stage 3/5. E/M/L2C: Early, mid late 2-cell stage; E/M/LB: Early, mid late blastocyst. ICM: inner cell mass. Scale bars equal 20 um in all panels. (D) Snapshots of a time-lapse movie of an embryo with a maternal *EsrrbTdT* knock-in allele and a paternal *H2b-Egfp* transgenic allele.

**Supplementary Information, Fig.S 2.**
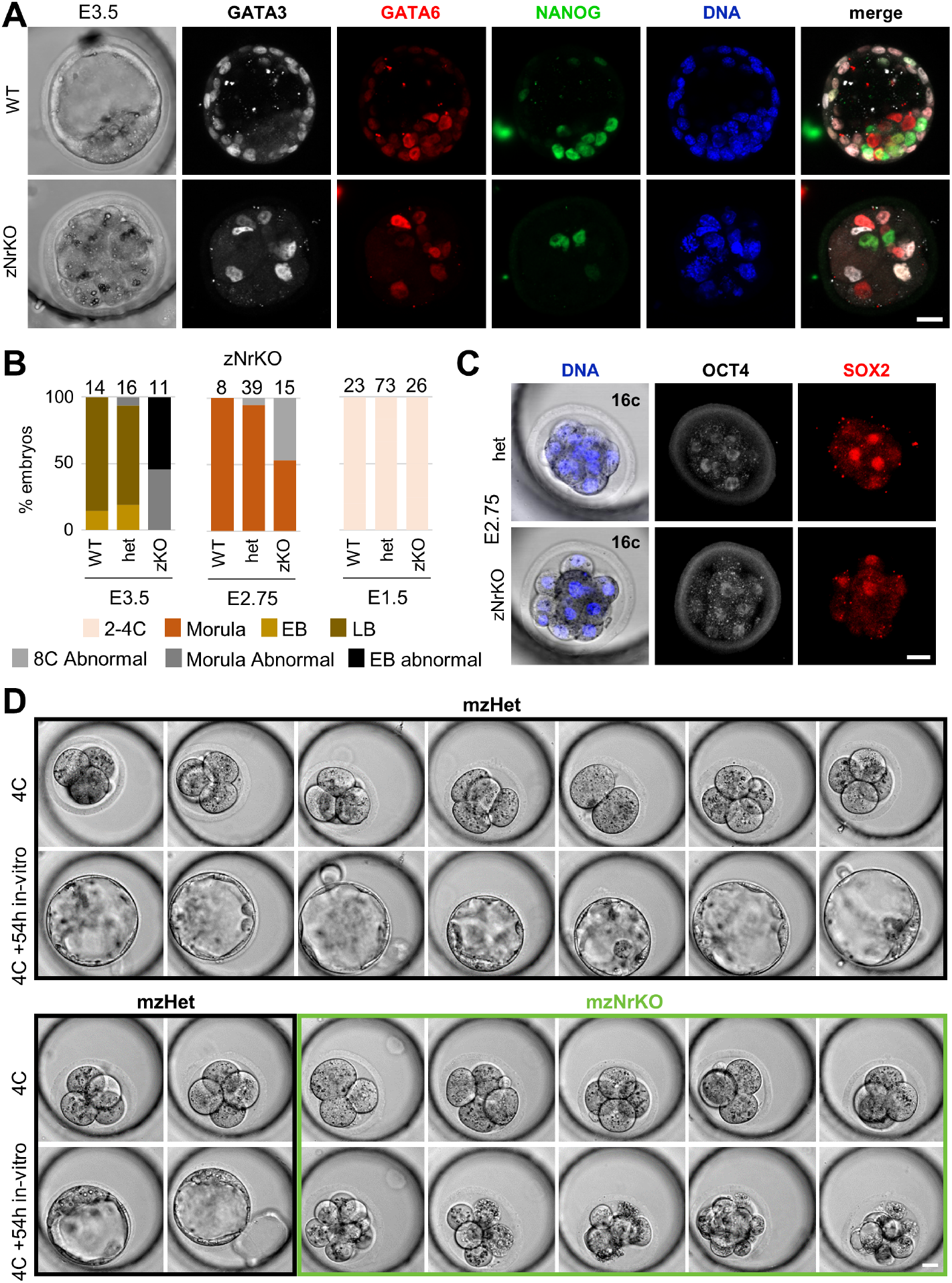
Compromised development of zygotic only *Nr5a2* mutants. (A) GATA3, GATA6 and NANOG immunostaining of zygotic *Nr5a2* heterozygous and KO embryos at E3.5. Scale bars equal 20 um in all panels. (B) Rate of development stages observed for each genotype in E1.5 to E3.5 litters from crosses generating zNrKO embryos. (C) OCT4 and SOX2 immunostaining of heterozygous and zNrKO littermate embryos at E2.75. (D) Pictures of all individual embryos from a litter issued from a cross between a *Zp3Cre*^*tg/0*^; *Nr5a2*^*flox/del*^ female and a *Nr5a2*^*del/+*^ male immediately after their recovery at late 2C/early 4C stage (hCG + 52h) and after 54 hours in culture (hCG + 106h).

**Supplementary Information, Fig.S 3.**
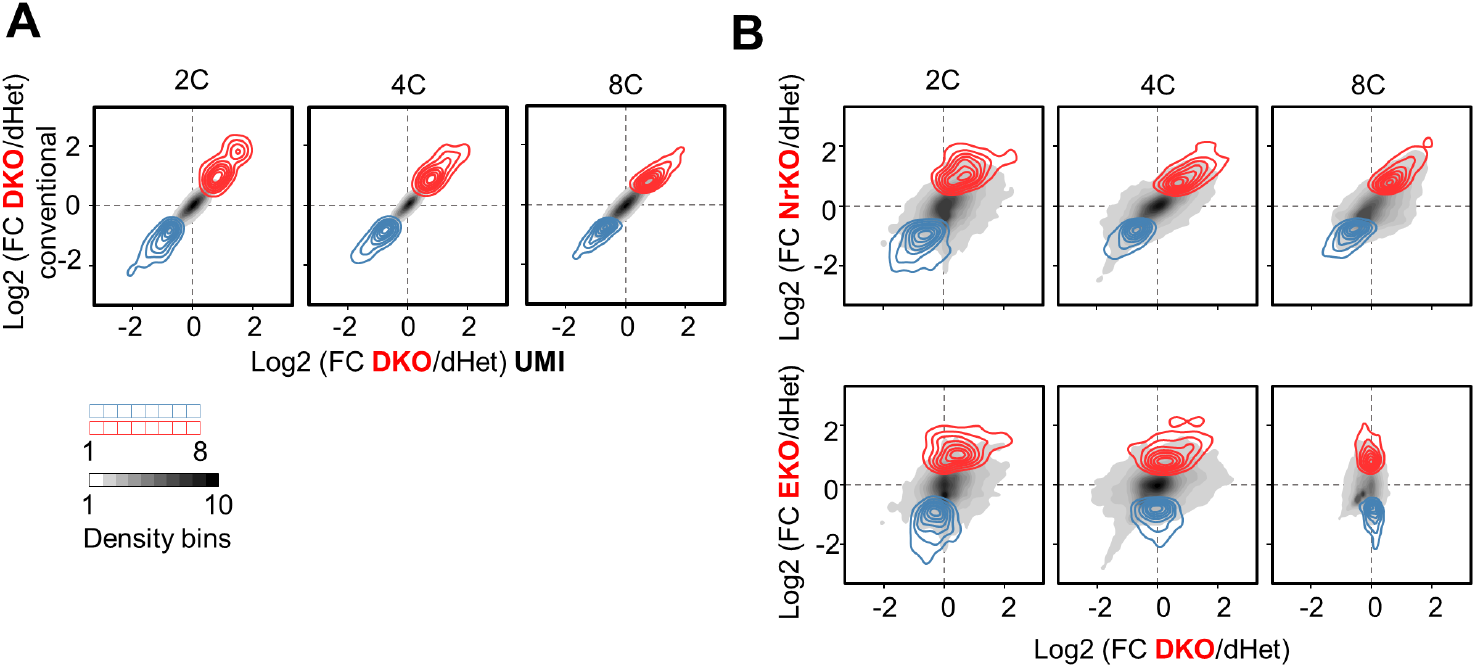
*Nr5a2* is dominant over *Esrrb* in 2-8C embryos. (A) Fold change in all expressed genes (TPM >10) in response to the inactivation of both *Nr5a2* and *Esrrb* (DKO) at different stages of development, as judged quantifying all reads, or relying on UMI counts. Red and blue distributions (divided in 8 bins) identify significantly upregulated or downregulated genes: absolute Fold change > 1.5; False discovery rate (FDR) < 0.1; cutoff 10 transcripts per million (TPM) - in DKO embryos at each stage, based on conventional analysis. The gray shades represent the density distribution of all genes, divided in 10 bins. (B) Fold change in expression of all genes in response to the single maternal and zygotic knockout of *Nr5a2* (NrKO) or *Esrrb* (EKO) versus the inactivation of both genes (DKO) at different stages of development. Red and blue distributions identify significantly upregulated or downregulated genes at each stage.

**Supplementary Information, Fig.S 4.**
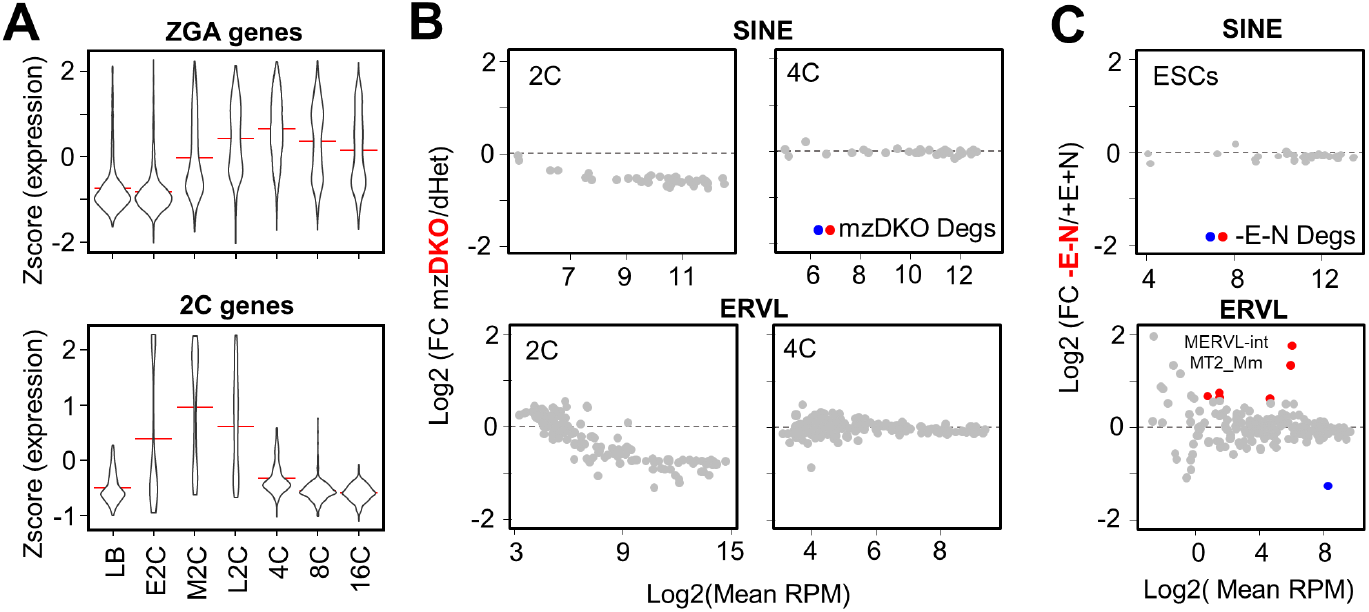
NR5A2/ESRRB repress expression of repeats linked to the 2C stage in ES cells, but not 2 or 4C embryos. (A) Relative expression levels **(18)** of 1984 genes activated during ZGA (as defined previously **(11)** and expressed in our datasets) or of 290 genes peaking at the 2C stage (see methods). (B,C) Fold change in expression of SINE or ERVL repeats in response to the maternal and zygotic KO of *Nr5a2* and *Esrrb*, at the 2 and 4C stage (B), or the loss of ESRRB/NR5A2 in ES cells **(12)** (C). Red and blue identify significantly upregulated or downregulated repeats in mzDKO embryos or DKO ES cells (absolut Fold change > 1.5; False discovery rate (FDR) < 0.1).

