## Supplemental Methods for "*Nr5a2* is essential for morula development"

**Festuccia et al.**

**Contents:**


**Supplementary Methods** **page 2**

1/ Animals and embryos production. page 2
*1a/ Animals husbandry.*
*1b/ Embryos production.*
*1c/ Embryo genotyping.*

2/ ESC culture conditions. page 3

3/ Imaging. page 4
*3e/ Embryo immunofluorescence.*
*3b/ Imaging of embryos.*

4/ Chromatin immuniprecipitation. page 5
*4b/ Chromatin preparation.*
*4c/ Chromatin immunoprecipitation (ChIP).*

5/ Single-embryo RNA-seq. page 6
*5b/ Library preparation.*
*5a/ SPRI Bead preparation.*

6/ Computational Methods. page 9
*6a/ Data availability.*
*6b/ Single-embryo RNA-seq analysis*.
*6c/ TF binding motif analysis.*
*6d/ Repetitive element expression.*
*6e/ Other computational analyses.*

**Supplementary references** **page 12**

### Supplementary Methods

#### *1/ Animals and embryos production*.

*1a/ Animals husbandry.*

All experiments were conducted according to the French and European regulations on care and protection of laboratory animals (EC Directive 86/609, French Law 2001-486 issued on June 6, 2001) and were approved by the Institute Pasteur ethics committee. Animals were kept on an inverted day/night cycle with 12 hours light cycle from 1 pm to 1 am.

*1b/ Embryos production.*

Embryos were staged according to the date of the vaginal plug (E0.5). Except for E4.25 (late blastocyst) embryos that were recovered from natural mating, embryos were collected from hormonally stimulated females following intraperitoneal injection of pregnant mare serum gonadotropin (PMS**G**) (2.5 IU/mouse, Chrono-gest ref A214A01 COSMO BIO), and 42-48h after with an additional injection of human chorionic gonadotropin (hCG) (5IU/mouse, Chorulon ref A242A01 COSMO BIO). Immediately after the second injection females were mated with appropriate males. Embryos were collected from females 20, 52, 65/75 or 95 hours after hCG in EmbryoMax FHM HEPES Buffered Medium w/o Phenol Red (SIGMA ALDRICH, Cat# MR-025D) for zygote, late E1.5 (late 2C/early 4C), E2.5 (8C-stage / morula stage) or E3.5 (blastocyst stage) respectively. mzKO embryos were collected from superovulated or natural mating *Zp3Cre*$^{tg/0}$*; Nr5a2*$^{flox/del}$ and/or *Esrrb*$^{flox/del}$ females, mated with *Nr5a2*$^{del/+}$ and/or *Esrrb*$^{del/+}$ males maintained on an essentially C57BL/6 genetic background. Zygotic *Nr5a2* KO embryos were recovered from crosses between *Nr5a2*$^{del/+}$ males and females maintained on a mixed C57BL/6/CD1 background. To visualize nuclei/chromatin during live imaging experiments, heterozygous *CAG::H2b-Egfp* (*1*) mice (on an essentially CD1 background) were crossed with *Esrrb*$^{TdT/+}$ mice.

*1c/ Embryo genotyping.*

Embryos were lysed in 5-15 µl of lysis buffer (10 mM Tris pH8 ; 50 mM KCl , 0.01 % gelatine + 300 µg/ml PK) at 56 °C for 1h followed by 10min at 95 °C in a thermocycler. 5 µl of lysate were used for 21 rounds of amplification using Phusion High-Fidelity DNA Polymerase (NEB, M0530L) in a total volume of 25 µl (13 µl H2O, 5 µl buffer 5X, 1.25 µl P1+P2+P3 (10 µM each), 0.5 µl dNTP (10 mM), 0.25 µl Phusion). 1 µl of the pre-amplified material was then used for 21-24 cycles of a similar nested PCR reaction, also in 25 µl. The amplification program for both steps was as follows:

30 sec 98°C
Followed by 21-24 cycles of:
15 sec 98°C
15 sec 64°C
1 min 72°C
Then:
3 min 72°C

Primers for 1st step *Esrrb*:
Esrrb del fw: CACGGTCAGCTTCCACTTTT
Esrrb flox fw: ACCATTCAAGGTGCGTGGTG
Esrrb rv: TGAGTCTTAGAGTTGAAATCCTTGT

Primers for 2nd step *Esrrb*:
Esrrb del fw: GGG TCT CTG ATT TGA AGT TTA CG
Esrrb flox fw: TATTAATTCCAAGTCTCGTTTCCTG
Esrrb rv: TATTAATTCCAAGTCTCGTTTCCTG

Primers for 1st step *Nr5a2*:
Nr5a2 del fw: CATAAGGGCTCAGTGGCAC
Nr5a2 flox fw: TAATAACTAAGAAGCAGAAAGCATGC
Nr5a2 rv: CTTCACTGGCTGCCAAGCTG

Primers for 2nd step *Nr5a2*:
Nr5a2 del fw: TACTGGTGATTTCCTGAGAGTACA
Nr5a2 flox fw: TAAGAAGCAGAAAGCATGCCAAG
Nr5a2 rv: CATTCTTCGGCAGTTGAGAGTGA

***2/ ESC culture conditions.***

ES cells were cultured on 0.1% gelatine (SIGMA, G1890-100G) in DMEM + GlutaMax-I (Gibco, 31966-021), 10% FCS (Gibco 10270-098), 100 µM 2-mercaptoethanol (Gibco, 31350-010), 1× MEM non-essential amino acids (Gibco, 1140-035) and 10 ng ml−1 recombinant LIF (MILTENYI BIOTEC, 130-099-895). Cells were passaged 1:10 every 2–3 days.

***3/ Imaging.***

*3e/ Embryo immunofluorescence.*

Embryos were fixed for 20 min at room temperature in 4% paraformaldehyde (Euromedex ref 15714) or on ice in 8% paraformaldehyde, 1X PhosSTOP for NR5A2 IF detection. Embryos were incubated in permeabilized/blocking solution in PBS with 0.1% Triton-X100 and 10% donkey serum (or 1% BSA for NR5A2 staining) for 30 minutes at room temperature. Embryos were treated with primary antibodies diluted in PBS with 0.1% Tween20 and 10% Donkey serum solution at 4°C overnight. After three washes with PBS with 0.1 % Tween20, embryos were incubated with secondary antibodies diluted 1/300 in PBS with 0.1% Tween20 for at least 1-2 hours. Hoechst 33342 (1.6 µM, Thermo Fisher) was included during incubation with secondary antibodies. The following antibodies were used: anti-NANOG (1:100; CosmoBio ref REC-RCAB0002PF), OCT4 (1:100; Santa Cruz ref sc5279), SOX2 (1:200; eBioscience, ref 14-9811-80), NR5A2 (1:100; R&D/Perseus ref PP-H2325-00), GATA6 (1:100; Cell Signaling Technology, ref 5851), GATA3 (1:100; R&D, ref AF2605), AlexaFluor 488 Donkey anti-mouse (1:300; Invitrogen, ref A21202) or anti-rat (1:300; Invitrogen, ref A21208), AlexaFluor 546 Donkey anti-rabbit (1:300; Thermo Fisher, ref A10040) and AlexaFluor 647 Donkey anti-goat (1:300; Thermo Fisher ref A21447).

3b/ *Imaging of embryos.*

All images were acquired by using a Zeiss LSM 900 Airyscan 2 Multiplex 4Y confocal microscope. Embryos were placed individually in wells of a customized microfabricated device (*2*) in PBS1X, 0.1 % Tween20. Images were acquired using a long working distance Plan-Apochromat objectives without immersion (20×/0.8), bi-directional scanning, 2x zoom and 1 airy unit pinhole. Stacks were created with a 2µm step, generating 16-bit at a confocal resolution images (1293x1293 Pixels).

For live imaging, embryos were cultured for 4 days in a 24-hour pre-equilibrated EmbryoMax KSOM medium (SIGMA ALDRICH, Cat# MR-121D) covered with FertiCult mineral oil (FertiPro, MINOIL050) and maintained in a humidified incubation chamber at 37°C in 8% $CO_2$. To minimize phototoxicity illumination for H2B-EGFP and ESRRB-TdT signal acquisition, low resolution images were generated with 512 x 512 pixels size, only 11 Z stack per embryos with a pinhole aperture of 100 µm imaged at a 15 min time interval.

***4/ Chromatin immuniprecipitation.***

*4b/ Chromatin preparation.*

*Fixation.* $10^7$ ES cells were crosslinked in 2 ml of freshly prepared PBS-DSG 2 mM at pH 7.0 (Sigma, 80424-5 mg) for 50 min at room temperature with occasional shaking. After pelleting and washing once in PBS, cells were incubated for 10 min in 2 ml PBS 1% formaldehyde (Thermo, 28908). Crosslinking was stopped with 0.125 mM glycine for 5 min at room temperature. Cells were pelleted and washed with ice-cold PBS. *Chromatin preparation*: Cells were resuspended in 2 ml of swelling buffer (25 mM Hepes pH 7.95, 10 mM KCl, 10 mM EDTA) freshly supplemented with 1× protease inhibitor cocktail (PIC-Roche, 04 693 116 001) and 0.5% NP-40. After 30 min on ice, the suspension was passed 40 times in a dounce (only for asynchronous populations). FCells were then centrifuged and resuspended in 300 µl of TSE150 (0.1% SDS, 1% Triton, 2 mM EDTA, 20 mM Tris-HCl pH8, 150 mM NaCl) buffer, freshly supplemented with 1× PIC. Samples were sonicated in 1.5 ml tubes (Diagenode) using a Bioruptor Pico (Diagenode) for 7 cycles divided into 30 s ON–30 s OFF sub-cycles at maximum power, in circulating ice-cold water. After centrifugation (30 min, full speed, 4 °C), the supernatant was stored at −80 °C. Five microlitres was used to quantify the chromatin concentration and check DNA size (typically 200–350 bp).

*4c/ Chromatin immunoprecipitation (ChIP).*

Chromatin from $2.10^6$ cells was used for each ChIP experiment. The chromatin was pre-cleared for 3 hours rotating on-wheel at 4 °C in 300 µl of TSE150 containing 50 µl of pG Sepharose beads (Sigma, P3296-5 ML) 50% slurry, previously blocked with BSA (500 µg ml$^{-1}$; Roche, 5931665103) and yeast tRNA (1 µg ml$^{-1}$; ThermoFisher Scientific, AM7119). Immunoprecipitations with anti-Flag mouse monoclonal (1 µg) (Sigma-Aldrich, F1804), anti-HA rabbit polyclonal (1 µg) (Abcam, Ab9110) were performed overnight rotating on-wheel at 4 °C in 500 µl of TSE150. Twenty microlitres was set apart for input DNA extraction and precipitation. Twenty-five microlitres of blocked pG beads 50% slurry was added for 4 h rotating on-wheel at 4 °C. Beads were pelleted and washed for 5 min rotating on-wheel at room temperature with 1 ml of buffer in the following order: 3 × TSE150, 1 × TSE500 (as TSE150 but 500 mM NaCl), 1× washing buffer (10 mM Tris-HCl pH8, 0.25M LiCl, 0.5% NP-40, 0.5% Na-deoxycholate, 1 mM EDTA), and 2 × TE (10 mM Tris-HCl pH8, 1 mM EDTA). Elution was performed in 100 µl of elution buffer (1% SDS, 10 mM EDTA, 50 mM Tris-HCl pH 8) for 15 min at 65 °C after vigorous vortexing. Eluates were collected after centrifugation and beads rinsed in 150 µl of TE-SDS1%. After centrifugation, the supernatant was pooled with the corresponding first eluate. For both immunoprecipitated and input chromatin, the crosslinking was reversed overnight at 65 °C, followed by proteinase K treatment, phenol/chloroform extraction and ethanol precipitation. Chromatin was resuspended in approximately 100µl of water and real-time RT–PCR reactions were performed in triplicate in 384-wells plates with a 480 LightCycler (Roche) using a LightCycler 480 SYBR Green I Master mix (Roche cat. 04887352001). Five microliters were used per reaction. Enrichment was calculated over inputs after accounting for the appropriate dilution factor and presented at bound and control genomic position. PCR primer sequences are listed below:

Subtelomeric region of Chr 13: fw CAGGGTTCAGGGGTCAAGGG; rv CCTACCCTCACCCTGTTCCTCA.

Control: fw GCCCAGGCAGGAGTAATTGG; rv GGCGTTCAGTAAGTTCCCCG.

***5/ Single-embryo RNA-seq.***

*5b/ Library preparation.*

The following protocol is derived from FLASH-seq (*3*), with two main modifications: 1/ Genotyping primers are added during the cDNA amplification step; 2/ A phosphate group is added to the 3' of the template switching primer to exclude the possibility it might prime DNA synthesis during the cDNA amplification step, which would result in tagging single transcripts with multiple UMI barcodes.

Embryos were isolated and directly transferred by mouth pipetting to PCR tubes in 5 µl of the following lysis solution:

- 0.1 µL Triton X-100 (10% v/v, Sigma-Aldrich, T8787)
- 1.2 µL dNTP mix (25 mM each, ThermoFisher Scientific, R1121)
- 0.46 µL 5'Bio-AAGCAGTGGTATCAACGCAGAGTAC$T_{30}$VN-3', 100 µM (supplied by IDT). Only polyadenylated transcript are detected by this protocol.
- 0.46 µL 5' Biotin-AGTGGTATCAACGCAGAGTAC G ATC NNNNNNNN rGrGrG-3'Phospate, 100 µM (supplied by IDT). AGTGGTATCAACGCAGAGTAC is a suppression PCR sequence. Random Ns form the UMI tag, and ATC is a 3bp barcode that can be varied if multiple samples are to be mixed before tagmentation, and only UMI reads are considered. rGrGrG are required for template switching. The 3' Phosphate inhibits priming in following steps.
- 0.15 µL recombinant RNase Inhibitor (40 U/µL, ThermoFisher Scientific, AM2684)

- 0.06 µL Dithiothreitol (DTT, 100 mM; part of the Superscript IV kit, ThermoFisher Scientific, 18091050)
- 1 µL betaine (5M, Sigma-Aldrich, 61962-50G)
- 0.45 µL dCTP (100 mM, Sigma-Aldrich, DNTP100)
- Water to 5 µL final.

Tubes were incubated 3 min at 72C in a thermocycler, and immediately transferred on ice. RT-PCR was performed by adding 20µl of the following mix to each sample:

- 1.19 µL Dithiothreitol (DTT, 100 mM; part of the Superscript IV kit, ThermoFisher Scientific, 18091050)
- 4 µL betaine (5M, Sigma-Aldrich, 61962-50G)
- 0.23 µL magnesium chloride (1 M, ThermoFisher Scientific, AM9530G)
- 0.48 µL recombinant RNase Inhibitor (40 U/µL, ThermoFisher Scientific, AM2684)
- 0.25 µL Superscript IV Reverse Transcriptase (200 U/µL, ThermoFisher Scientific, 18091050)
- 12.5 µL KAPA HiFi Hot-Start ReadyMix (2x, Roche; cat. KK2502)
- Water to a final volume of 20 µL.

Samples were incubated in a thermocycler set to execute the following programme:

Hold at 50°C, and after placing tubes skip to
60 min at 50°C
3 min at 90°C
Hold at 4°C

Tubes were transferred on ice and 0.5µl of a 100 µM solution of amplification primer (TCGTCGGCAGCGTCAGATGTGTATAAGAGACAG AAGCAGTGGTATCAACGCAGAGTAC G) were added to each sample. 1µl of a solution of genotyping primers (see embryo genotyping section) at 10µM each was also added in each tube.

Tubes were then transferred to a thermocycler, set to the following programme:

Hold at 4°C and after placing tubes skip to:
20 sec 98°C
20 sec 64°C
6 min 72°C

Followed by 20 cycles of:

20 sec 98°C
20 sec 67°C

6 min 72°C, adding 12 seconds per cycle
Hold at 4°C

Amplification products were purified using 1x volume of SPRI magnetic beads (homemade, see the SPRI bead preparation and use section), eluting in 15 µl water. 1 µl was set aside to perform the second step of the nested PCR genotyping protocol (see embryo genotyping section).
Fragment size distribution was checked on a Genomic DNA Screentpape (Agilent Technologies, 5067-5365/6), with a typical lengths ranging from less than 1 kb to more than 10 kb.
For tagmentation, each sample was diluted to 150 pg/µl. 2 µl of diluted cDNA were mixed with 2µl of Amplicon Tagment Mix (ATM) and 4 µL of Tagment DNA Buffer (TD) from an Nextera XT DNA Library Preparation Kit (Illumina, FC-131-1024), and incubated for 8 min at 55°C, in a thermocycler. Tn5 was released from DNA by adding 2 µL of Neutralization Buffer (NT), followed by a 5 min RT.
Enrichment of UMI containing fragments in the tagmented DNA was performed by adding 5 µL of Nextera PCR Master Mix (NPM) and 1 µl of 5 µM amplification primer (TCGTCGGCAGCGTCAGATGTGTATAAGAGACAG AAGCAGTGGTATCAACGCAGAGTAC G, as before). KAPA HiFi Hot-Start ReadyMix (2x, Roche; cat. KK2502) can substitute the Nextera PCR Master Mix, adapting volumes as required. Tubes were incubated as follows:
72°C for 3 min
95°C for 30 sec

Followed by 10 cycles of linear enrichment PCR (or more depending on how many UMI reads are required):

95°C for 10 sec
55°C for 30 sec
72°C for 30 sec
Hold at 4°C

4 µl of a 2.5 µM solution of each of the following amplification primers were added on ice:
Universal primer
AATGATACGGCGACCACCGAGATCTACACTCGTCGGCAGCGTCAGATGTGTATAAGAGACAG
Indexed primer
CAAGCAGAAGACGGCATACGAGATNNNNNNNNGTCTCGTGGGCTCGGAGATGTGTATAAGAGACAG. The 8 Ns are to be replaced with the Illumina index sequence of choice.

Tubes were returned to the thermocycler held at 4°C and then subjected to 14 iterations of the cycle:

95°C for 10 sec
55°C for 30 sec
72°C for 30 sec
Followed by:
72°C for 5 min
4°C hold

Amplification products were purified using 1-1.4x volume of SPRI magnetic beads, eluting in 30 µl water. Fragment size distribution was checked on a DNA Hi Sensitivity D1000 HS Screentape (Agilent Technologies, 5067-5584/5).

*5b/ SPRI Bead preparation.*

Preparation: 1 ml Sera-Mag™ Magnetic SpeedBeads™, carboxylated, 1 µm, 3 EDAC/PA5 (GE Healthcare Life Sciences, Cat #65152105050250) were washed 3 times with a TE-Tween solution (10 mM Tris HCl pH 8, 1 mM EDTA, 0.05% Tween 20, pH 8.0) and resuspended in TE-Tween-20% PEG 8000 solution (10 mM Tris HCl pH 8, 1 mM EDTA, 0.05% Tween 20, pH 8.0). DNA purification: 1 or 1.2x sample volumes of SPRI beads were added to each tube and samples transferred to a 96 well plate. After incubating for 5 min, the plate was put on a 96S Super Ring Magnet (Alpaqua, A001322), beads were allowed to separate completely, and the supernatant removed without disrupting the bead pellet. Beads were washed twice with approximately 100 µl of 70% Ethanol and the supernatant completely removed. DNA was eluted in water after separating the beads one last time.

**6/ Computational Methods.**

*6a/ Data availability.*

Samples are summarised in Table S5, which also indicates preparation day, embryonic stage, and genotype. Fastq files from the listed samples, generated by trimming and splitting UMI and non-UMI reads, will be deposited in the GEO database.

*6b/ Single-embryo RNA-seq Analysis*.

Single-end sequences (50-85bp, using 50bp sequencing kits) were generated using an Illumina Next-seq 500 or 2000 instruments. After trimming Illumina adapter sequences with cutadapt (*4*) using options -b CTGTCTCTTATA --minimum-length=1, reads were split in UMI containing or not with Julia (*5*), trimming 11bp after the sequence "AAGCAGTGGTATCAACGCAGAGTACGATC". For UMI, reads were further processed in Julia to remove the invariant adapter sequence, extract the identifier, and the UMI/ transcript sequence pairs used to remove duplicates. Typically UMI libraries displayed 30% of unique reads. Note that a much lower duplication rate is to be expected while analysing conventional reads, as in our protocol UMIs are anchored to the 5' of transcripts, and thus reads are not randomly distributed and less diverse. Conventional or UMI reads were then aligned to the mm10 genome using STAR (*6*) and quantified by RSEM (*7*) through the RSEM-STAR pipeline, using gencode.v25 annotations and additional options "--seed 1618 --calc-pme --calc-ci --estimate-rspd --forward-prob 0.5" for conventional reads and "--seed 1618 --calc-pme --calc-ci --estimate-rspd --forward-prob 1" for UMI reads. RSEM estimated read counts per sample were rounded for use with DESeq2 (*8*). For all differential expression tests DESeq2 was run without independent filtering and without any fold change shrinkage. Genes were considered as differentially expressed if their mean expression exceeded 10 TPM in at least one genotype for a given stage, and using a <> 1.5 fold change and <0.1 FDR cutoff.

*6c/ TF binding motif analysis.*

First, fastq files were obtained from the GEO series GSE66581 (*9*). Reads were aligned with Bowtie2 (*10*) to the mm10 genome, with options "-k 1". Experimental replicates for each stage were merged and peaks were called using MACS2 (*11*), which defined accessible regions. To discover transcription factor binding motifs at these accessible regions in the proximity of differentially expressed genes, coordinates of all transcript annotated in the mm10 genome were obtained from ENSEMBL using the biomaRt package (*12*) (dataset = "mmusculus_gene_ensembl", host="apr2020.archive.ensembl.org"). For each gene, and separately at embryonic each stage, the transcript showing maximal expression was selected by quantifying the UMI read counts in a window of -50/+150 bp from the TSS using the bamsignals R package (*13*). Only genes showing counts greater than zero for at least one isoform were considered. Separately for each stage, we identified the sequences of the accessible regions falling in a window of +/- 20Kb from the TSS of each expressed gene. For de-novo motif discovery sequences were analysed with RSAT (*14*) peak motif algorithm, with the following command options:
"-v 1 -max_seq_len 1000 -markov auto -disco oligos,dyads,positions,local_words -nmotifs 10 -minol 6 -maxol 8 -merge_lengths -2str -origin center -motif_db jaspar_core_nonredundant _vertebrates tf $RSAT/public_html/motif_databases/JASPAR/Jaspar_2020/nonredundant/ JASPAR2020_CORE_vertebrates_non-redundant_pfms.tf –task,purge,seqlen,composition,

disco,merge_motifs,split_motifs,motifs_vs_motifs,timelog,archive,synthesis,small_summary,motifs_vs_db -prefix peak-motifs -noov -img_format png”

To identify occurrence of Nr5a2/Esrrb or SINE Y5 the relative pfm or pwm matrixes were obtained using the Jaspar2014 (*15*) R package (searching “Nr5a2” matrix by name; or MA0505.1), and from (*16*). After trimming, the same set of sequences from accessible region used for de-novo discovery was also scanned for occurrence of Nr5a2 or SINE Y5 motifs using the TFBT R package (*17*), setting a quality threshold of > 85%. Only matches displaying 0-3 mismatches to the consensus were retained for SINE. For Nr5a2/Esrrb motifs with 0-1 mismatches, in addition to the allowed T or C variation of the 7th base, were considered. The highest scoring matches to the motifs was retained for each region.

*6d/ Repetitive element expression.*

Non-UMI reads were were mapped to the mm9 genome using bowtie2, and split by the RepEnrich2_subset command of RepEnrich2 (*18*), using a mapq value of 30 to subset uniquely mapping from multi-mapping reads. Repeat expression was then quantified by RepEnrich2 using the default repeatmasker file mm9.fa.out.gz downloaded from repeatmasker.org. Differential enrichment analysis was performed using a generalized linear model in EdgeR (*19*), following a standard procedure (https://github.com/nerettilab/RepEnrich2). Repeats were called as differentially expressed using a <> 1.5 fold change and <0.1 FDR cutoff.

*6e/ Other computational analyses.*

Gene ontology analysis was performed pooling all differentially express genes, and using the enrichGO function of the ClusterProfiler R package (*20*), with options “ont ="BP", pAdjustMethod = "BH" (Bonferroni-Holm correction for multiple comparisons), minGSSize = 10, maxGSSize = 170 (exclude terms associated with less than 10 and more than 170 genes), pvalueCutoff = 0.01”. Results plotted with the command emapplot, after calculating the pairwise term similarities and displaying the first 150 enriched terms. Dot violin and box plots were generated in R using ggplot2 (*21*). Venn diagrams were made with eulerr (*22*), heatmaps were made with ComplexHeatmap (*23*). ChIP-seq profiles at telomeric regions were generated re-aligning datasets (GSM4605754, GSM4605774, GSM4605794, GSM6797321, GSM6797322, GSM6797323) from (*24, 25*) using bowtie with param --sensitive-local. Bigwig files were created from the alignments by the bamCoverage command of deeptools (*26*) with --binSize 20, and graphs generated from Bigwig using rtracklayer (*27*), smoothed with zoo (*28*) and plotted using the Gviz (*29*) and GenomicFeatures (*30*) R packages. PCA plots were generated in R considering genes with expression greater than 10 TPM in at least one sample, using the prcomp command, with options center = T. Tables reporting gene expression along pre-implantation murine development were obtained from GSE45719 (*31*). Genes upregulated during ZGA were selected based on this dataset, to have an expression >10 RPKM in late 2C or 4C embryos, and a fold change between early 2C and 4C stages > 4. Genes were further filtered to have mean expression greater than 0 TPM in all 2C-stage samples, and greater than 10 TPM for at least one genotype, in our study. Also using data from (*31*), "2C-specific" genes were selected to have expression greater than 5 RPKM at the early, mid or late 2C-stage, and so that their relative expression, setting max to 1 and min to 0, was <0.5 at 4C and <0.3 at all other stages. Genes were further filtered to have mean expression greater than 0 TPM in all 2C-stage samples, and greater than 10 TPM for at least one genotype, in our study.
