## Supplementary material for "*Nr5a2* is essential for morula development": Table S1

**Table S1: Genotypes of live progeny and embryos recovered at different stages of development, after the maternal or maternal and zygotic deletion of Nr5a2/Esrrb.**

***A/ Transmission of deleted Nr5a2 and Esrrb alleles to the live progeny of Zp3Cre^tg/0^*; *Esrrb^del/flox^* or *Nr5a2^del/flox^* females occurs at the expected frequency**

Genotype of live pups from females *Zp3Cre^tg/0^*; *Esrrb^del/flox^* females crossed with wild-type CD1 males

|  | transmitted allele | | | | |
| --- | --- | --- | --- | --- | --- |
| female genotype | *wt* | *flox* | *del* | *ND* | Total |
| Zp3Cre ^tg/0^; Esrrb^flox/+^ | 23 | 0 | 32 | 3 | 58 |
| Zp3Cre ^tg/0^; Esrrb^flox/del^ | 0 | 0 | 72 | 0 | 72 |

n=4 females per genotype

Genotype of live pups from females *Zp3Cre^tg/0^*; *Nr5a2^del/flox^* females crossed with wild-type CD1 males

|  | transmitted allele | | | | |
| --- | --- | --- | --- | --- | --- |
| female genotype | *wt* | *flox* | *del* | *ND* | Total |
| Zp3Cre ^tg/0^; Nr5a2 ^flox/+^ | 10 | 0 | 17 | 0 | 27 |
| Zp3Cre ^tg/0;^ Nr5a2 ^flox/del^ | 0 | 0 | 62 | 1 | 63 |

n= 3 control and n=4 mutant females respectively

***B/ mzKO o zKO embryos are recovered at the expected frequencies***

**mzEsrrb KO**

Genotype of embryos from *Zp3Cre^tg/0^*; *Esrrb^del/flox^* females crossed with *Esrrb^del/+^* males

|  | Number of embryos per genotype | | | |
| --- | --- | --- | --- | --- |
| Stage | *Esrrb^del/+^* | *Esrrb^del/del^* | *ND* | Total |
| E3.0 | *3* | *4* | *0* | 7 |
| E 3.5 + 20h | 6 | 11 | 0 | 17 |
| E 4.0 | 6 | 6 | 0 | 12 |
| Total | 15 | 21 | 0 | 36 |

n=1 female at E3.0; n=2 females at E3.5 + 20h and n=3 females at E4.0

**mzNr5a2 KO**

Genotypes of embryos from female *Zp3Cre^tg/0^*; *Nr5a2^del/flox^* females crossed with *Nr5a2^del/+^* males

|  | Number of embryos per genotype | | | |
| --- | --- | --- | --- | --- |
| Stage | *Nr5a2^del/+^* | *Nr5a2^del/del^* | *ND* | *Total* |
| E 2.5 | 19 | 14 | 3 | 36 |
| E 3.5 | 12 | 15 | 1 | 16 |
| E 4.0 | 16 | 14 | 0 | 30 |
| Total | 47 | 43 | 4 | 72 |

n=4 females at E2.5; n=2 females at E3.5 and n=3 females at E4.0

**zNr5a2 KO**

Genotypes of embryos from *Nr5a2^del/+^* females crossed with *Nr5a2^del/+^* males

|  | Number of embros per genotype | | | |  |
| --- | --- | --- | --- | --- | --- |
| Stage | *Nr5a2^+/+^* | *Nr5a2^del/+^* | *Nr5a2^del/del^* | *ND* | *Total* |
| E1.5 | 5 | 6 | 4 | 2 | *17* |
| E 2,5 | *8* | *39* | *15* | *8* | *70* |
| E 3.5 | 14 | 16 | 11 | 0 | 41 |
| Total | 27 | 61 | 30 | 10 | 128 |

n=1 female at E1.5; n=5 females at E2.5; n=3 females at E3.5

**mzDKO**

Genotypes of embryos from female *Zp3Cre^tg/0^*; *Esrrb^del/flox^*; *Nr5a2^del/flox^* females crossed with *Esrrb^del/+^; Nr5a2^del/+^* males

|  | Number of embros per genotype | | | | |  |
| --- | --- | --- | --- | --- | --- | --- |
| Stage | **DKO**  *Esrrb^del/del^*  *Nr5a2^del/del^* | **EKO**  *Esrrb^del/del^ Nr5a2^del/+^* | **NrKO**  *Esrrb^del+l^ Nr5a2^del/del^* | **DHet**  *Esrrb^del/+^ Nr5a2^del/+^* | *ND* | *Total* |
| E1.5 | 24 | 19 | 20 | 28 | 13 | 104 |
| E 2,5 | 23 | 29 | 23 | 37 | 4 | 116 |
| Total | 47 | 48 | 43 | 65 | 17 | 220 |

n=7 female at E1.5; n=6 females at E2.5
